# Ancient origins of complex neuronal genes

**DOI:** 10.1101/2023.03.28.534655

**Authors:** Matthew J. McCoy, Andrew Z. Fire

## Abstract

How nervous systems evolved is a central question in biology. An increasing diversity of synaptic proteins is thought to play a central role in the formation of specific synapses leading to nervous system complexity. The largest animal genes, often spanning millions of base pairs, are known to be enriched for expression in neurons at synapses and are frequently mutated or misregulated in neurological disorders and diseases. While many of these genes have been studied independently in the context of nervous system evolution and disease, general principles underlying their parallel evolution remain unknown. To investigate this, we directly compared orthologous gene sizes across eukaryotes. By comparing relative gene sizes within organisms, we identified a distinct class of large genes with origins predating the diversification of animals and in many cases the emergence of dedicated neuronal cell types. We traced this class of ancient large genes through evolution and found orthologs of the large synaptic genes driving the immense complexity of metazoan nervous systems, including in humans and cephalopods. Moreover, we found that while these genes are evolving under strong purifying selection as demonstrated by low dN/dS scores, they have simultaneously grown larger and gained the most isoforms in animals. This work provides a new lens through which to view this distinctive class of large and multi-isoform genes and demonstrates how intrinsic genomic properties, such as gene length, can provide flexibility in molecular evolution and allow groups of genes and their host organisms to evolve toward complexity.

## Introduction

Gene size varies among organisms and can change due to the addition of domains to proteins with increasing complexity^1^. However, while protein sizes remain consistent among eukaryotes^2^, absolute gene sizes within and among species can vary greatly^3–7^. The majority of these differences are caused by expansions of non-coding DNA, specifically within introns^3,5^. Average intron sizes are correlated with genome size^8,9^ and can impact a range of ecological and cellular processes^10,11^.

The consequences of gene size variation are only beginning to be understood. Many of the largest animal genes are commonly expressed in nervous tissue^7,12–15^, and are frequently mutated or misregulated in human conditions such as autism spectrum disorders^12^ and Rett syndrome^13^. Other studies have found that larger genes are less likely to undergo full duplication and more likely to exhibit alternative splicing^6,16^. However, the mechanisms underlying the acquisition of large, complex genes during evolution and their breakdown in disease are not yet fully understood.

Recent tools allow concrete orthology assignments of orthologous genes between diverse species^17,18^. This development provides the opportunity to move from averages to individual trajectories of gene and protein size during evolution. Here, we compare the size, age, and architecture of animal genes to provide insight into the origins of molecular diversity and complexity in many animals and their nervous systems.

## Results

### Relative gene size is preserved among species

To determine the evolutionary origins of large neuronal genes, we set out to define and characterize this set of genes across diverse species. This required first identifying orthologs and investigating the variation of gene architectural features.

In this work we use several terms to describe aspects of size associated with gene expression patterns and function. The term *gene size* refers to the length from the start of the first annotated exon in the genome to the end of the last annotated exon, including introns. This definition excludes 5’ and 3’ untranslated regions, because these are often under-annotated^3^. We measure and compare size in two ways: the *absolute size* and the *relative size*. The term *absolute size* refers to the number of base pairs. The term *relative size* refers to the ranked size relative to other genes within the same genome. We use the term *CDS size* to refer to the span of nucleotides within a mature RNA transcript that will eventually be translated into protein, which excludes introns and untranslated regions. *Protein size* is measured by the number of amino acids. These definitions are important because we will argue that both relative and absolute gene size contribute distinct aspects to the evolution of gene expression patterns and function.

The ratio of introns to intergenic sequences is nearly 1:1 in numerous model animals^3^. Hence, larger animal genomes typically have larger intronic content and thus larger genes^3,19^. Together with previous studies showing that orthologous proteins are encoded by genes with similar-sized CDS^2^, this would suggest that changes in gene sizes are a simple function of changes in genome size. While previous studies have compared aggregate measures of gene size or coding and non-coding DNA in different species^3,5,20^, we required gene-by-gene comparisons to investigate gene size variation during evolution and its impact on co-expression patterns and gene architecture. Therefore, we first set out to compare orthologous gene sizes among diverse eukaryotes.

We asked whether gene sizes in one species covary with orthologous gene sizes in distantly related species. We addressed this question by comparing rank orders of gene size between species. We focused our analysis on several diverse eukaryotes with chromosome-level genome assemblies in part because gene annotation quality is related to genome assembly completeness^3^. For this analysis, we identified one-to-one orthologs using OrthoFinder^18^, a highly accurate orthology inference tool that accounts for gene-length bias in detecting orthologs^17^. Despite orders-of-magnitude variation in absolute gene size, we found that relative gene size is largely maintained across species (**Fig. 1A,B, Extended Data Fig. 1**). This is true not only among vertebrates, which typically have significantly larger genes than invertebrates, but also in comparisons with cephalopods (see *Octopus sinensis* in **Fig. 1A,B**), which are of particular interest due in part to their evolution of large and complex nervous systems independent of vertebrates^21^. For the purpose of juxtaposition with gene sizes, **Figure 1C** displays protein sizes, which have previously been found to be nearly invariant among eukaryotes^2^. These results support the hypothesis that gene sizes are shifting together at the macroevolutionary scale. This also indicates that the largest genes in one species are among the largest in distantly related species, but can vary in absolute size by orders of magnitude.

**Figure 1.**
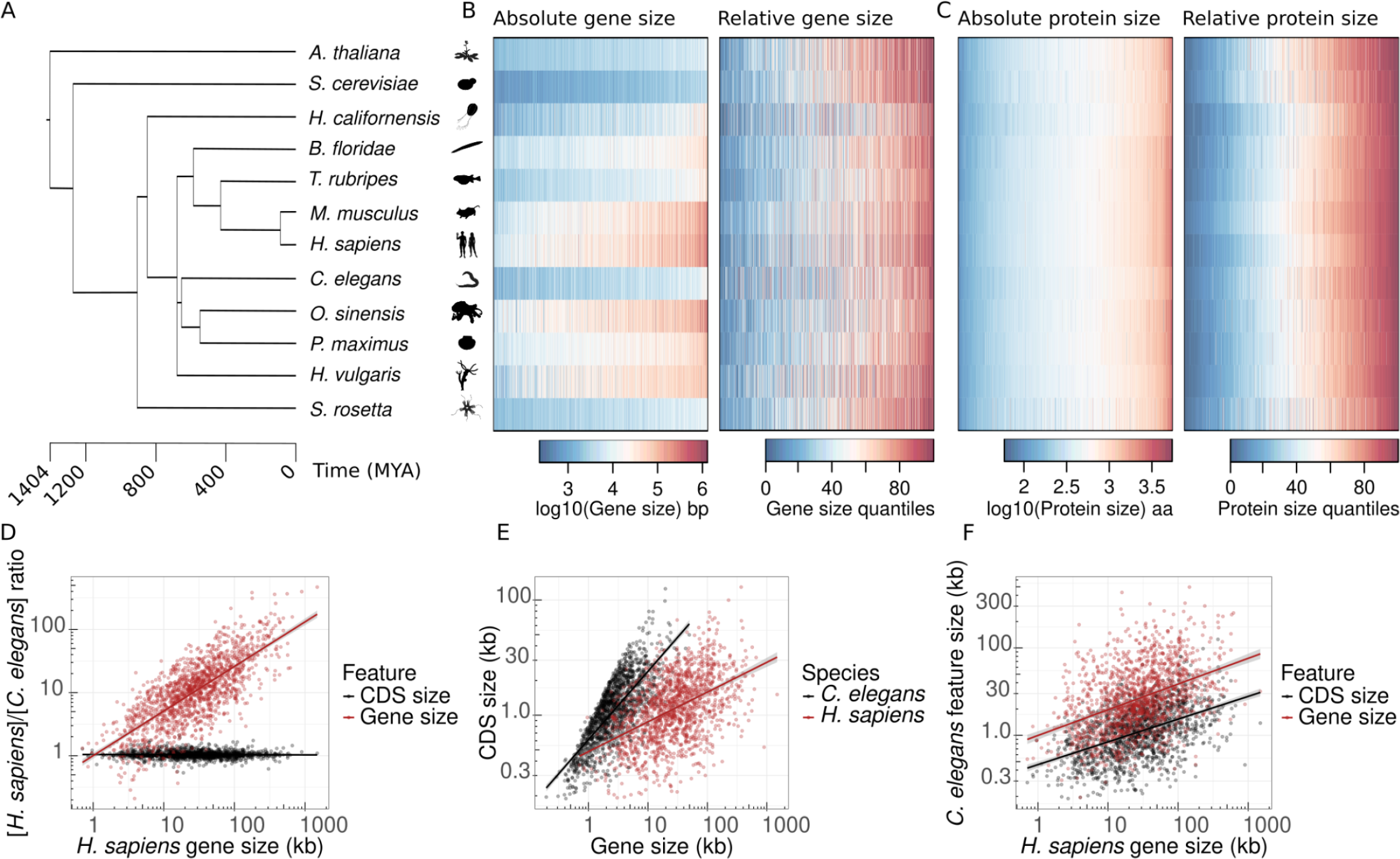
Absolute gene sizes vary by orders of magnitude among diverse species while relative gene sizes are maintained. (**A**) Phylogenetic tree of eukaryotic species with chromosome-level assemblies (excepting *S. rosetta*) used in this study. Branch lengths determined by TimeTree.org^23^. (**B**) OrthoFinder one-to-one ortholog gene sizes across (Left) Heatmap of absolute gene size (log10 bp) with genes (columns) ordered by the average gene size. (Right) Heatmap of relative gene size (gene size quantiles), with each ortholog binned into 100 quantiles to show the size ranking for the same gene across orthologs in different species. Genes (columns) are ordered by gene size quantile across all species. (**C**) (Left) Heatmap of absolute protein size (log10 aa) with proteins (columns) ordered by average protein size. (Right) Heatmap of relative protein size (protein size quantiles), with each ortholog binned into 100 quantiles to show the size ranking for the same proteins across orthologs in different species. Proteins (columns) are ordered by protein size quantile across all species. (**D-F**) Gene and CDS size of Ensembl one-to-one, high-confidence orthologs between *Homo sapiens* and *Caenorhabditis elegans*. Solid lines show linear models with 95% confidence intervals as ribbons. (**D**) CDS size remains relatively invariant, while gene size varies substantially. Ratios of [*H. sapiens*]/[*C. elegans*] gene and CDS size. (**E**) *C. elegans* gene and CDS size are both strongly correlated with orthologous gene sizes in *H. sapiens*. (**F**) Gene size is correlated with CDS size within individual genomes.

We sought additional evidence of the relationship between gene and CDS size (and hence protein size) by comparing one-to-one orthologs (obtained from Ensembl^22^) in *Homo sapiens* and the invertebrate nematode, *Caenorhabditis elegans*, which have some of the best characterized animal genomes. Humans shared a common ancestor with nematodes roughly 680 million years ago^23^, and since then the size of our haploid genome has expanded to more than 3 billion base pairs, roughly 30 times the size of the *C. elegans* haploid genome at around 100 million base pairs^24^. We found that while the CDS size of each orthologous gene is nearly invariant between species, the largest human genes can be more than 100 times the size of their orthologs in *C. elegans* (**Fig. 1D**). We also found that within *H. sapiens* and *C. elegans* genomes, CDS size is strongly correlated with gene size (**Fig. 1E**). When we compared the correlation of gene size in humans with either CDS size or gene size in *C. elegans*, we found these relationships to be similarly strong, suggesting that protein size and gene size are closely related on a macroevolutionary scale (**Fig. 1F**). These results are consistent with the known conservation of orthologous protein sizes among diverse eukaryotes^2^, while highlighting significant differences in absolute gene size that may underlie important aspects of gene function and expression.

### Specific neuronal functions enriched for large genes

One unusual feature of nervous tissue is the high number of genes with tissue-specific expression^25^. Previous studies observed that many of the largest genes are enriched for expression in the brain^7,12,13,15,26–28^. Using tissue-enriched genes provided by the Human Protein Atlas^25^, we quantified the number of tissue-enriched genes at each gene size and found more brain-enriched genes in the top 10% largest genes than in any other size range (**Fig. 2A**). This result is juxtaposed with the high number of small genes enriched for expression in testis and skin (**Fig. 2A**).

**Figure 2.**
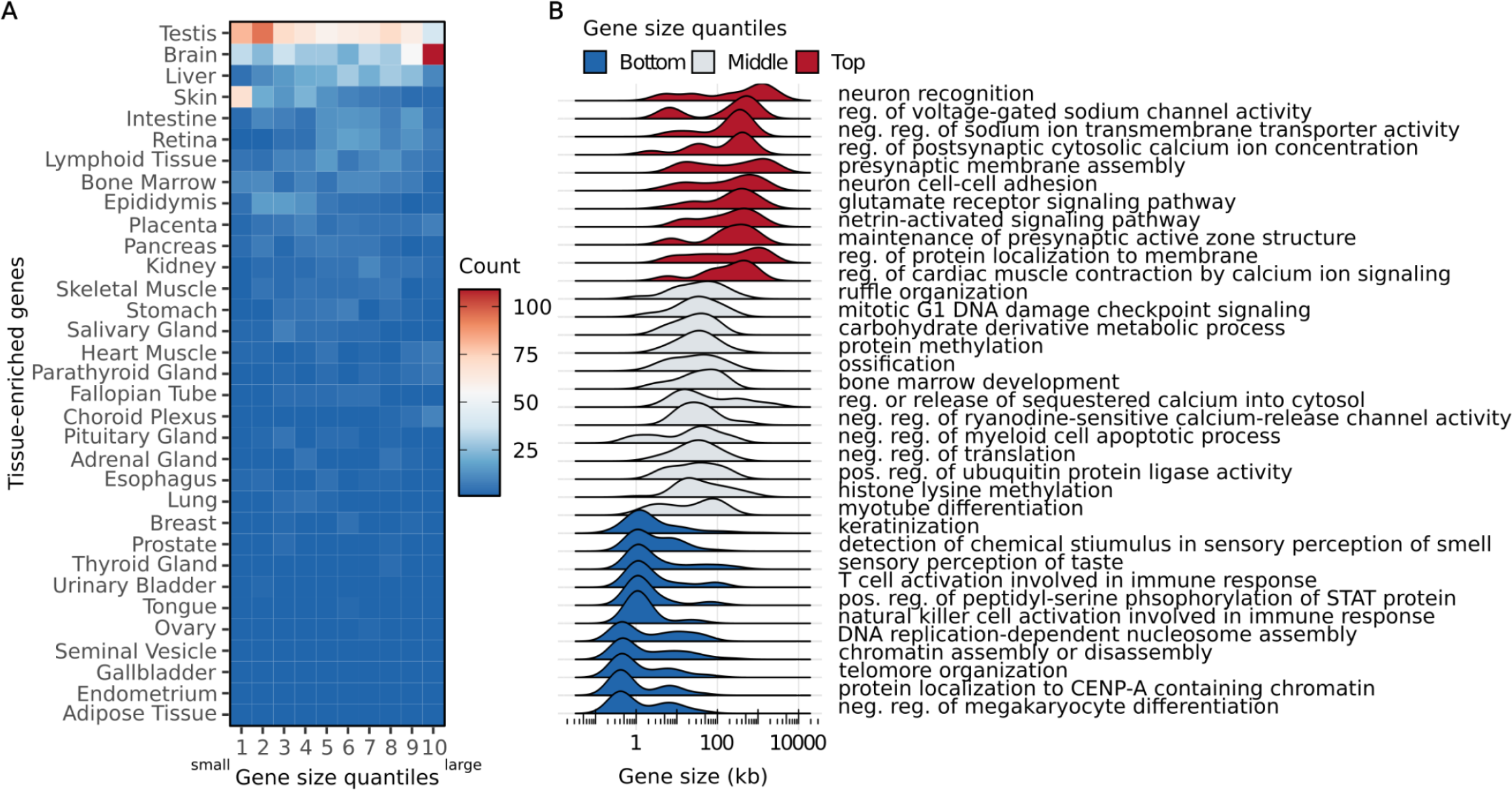
Brain tissue and many neuronal processes are enriched for large genes. (**A**) Heatmap of Human Protein Atlas tissue-enriched genes binned by gene size quantiles (10 bins). Heat colors show the number of genes in each bin. Tissues are ordered by the total number of enriched genes across all gene sizes in each tissue. (**B**) Density plots (joy plots) showing human GO biological terms filtered to display terms with the lowest deviation from median gene sizes (top 10, middle 10, bottom 10).

Previous studies have also shown that the largest genes are enriched for gene ontology (GO) terms associated with synaptic function^13^. We examined gene size distributions for GO terms associated with individual functions, which provided a striking picture in which some functions were associated with a majority of genes in a specific size class (**Fig. 2B**). In particular, many GO terms composed mainly of large genes are involved in neuronal function (e.g. neuron recognition, presynaptic membrane assembly, neuron cell-cell adhesion, etc.)(**Fig. 2B**). These results suggest there are classes of genes whose functions may benefit from (i) small condensed gene sizes, such as highly expressed genes^29–31^ and genes involved in rapid stress response^32^, or (ii) expanded gene sizes, such as neuronal genes with numerous isoforms. There may also be a third class of genes that (iii) do not benefit from either small or expanded gene sizes, or whose gene sizes are determined by currently unknown forces.

### Most large neuronal genes are ancient

Previous studies found that older genes on average are larger, experience stronger purifying selection, and evolve more slowly than younger genes^33–35^. However, these aggregate measures obscure certain features, such as the fact that many short ancient genes are evolving under strong purifying selection (e.g. histone genes^36^). We therefore sought a more detailed analysis on genes of specific ages and sizes.

Our analysis in **Figure 1** focused on genes with orthologs across diverse eukaryotes, and thus was necessarily limited to ancient conserved genes. To address whether most large genes are ancient and conserved, we used estimates of gene age based on the phylogenetic distribution of orthologs as described by Tong et al. (2021)^37^. We found that most of the larger protein-coding genes are indeed ancient, with the top 10% largest human genes averaging an inferred age of 953 million years old, whereas the top 10% shortest genes have an average inferred age of 62 million years old (**Fig. 3A,B**). We also found that compared with shorter genes the top 10% largest genes in our analysis have lower dN/dS scores between human and mouse (**Fig. 3C**). Furthermore, the largest genes also have higher gene order conservation (GOC)(**Fig. 3C**). These features indicate large genes are under stronger purifying selection and have similar local gene neighborhoods, which together suggest that large genes are highly conserved.

**Figure 3.**
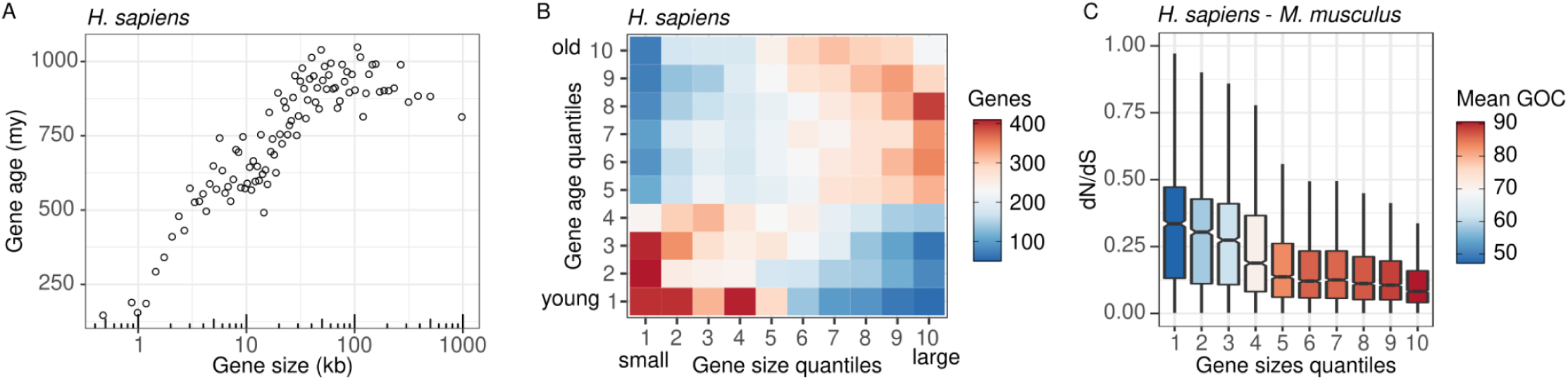
Most large genes are ancient, while most young genes are small. (**A**) Scatter plot of mean gene size versus mean gene age (my = million years) of genes binned by size (100 bins) in *H. sapiens*. (**B**) Heatmap of gene size quantiles versus gene age quantiles in *H. sapiens*. Heat colors show the number of genes within each tile and are capped between 50 and 410 genes. (**C**) The largest genes have lower dN/dS scores and higher gene order conservation (microsynteny) than short genes. Boxplots of *H. sapiens* vs. *M. musculus* dN/dS scores (from Ensembl one-to-one orthologs) across *H. sapiens* genes binned into 10 size quantiles. Heat colors show the mean gene order conservation (GOC).

Starting from a list of large brain-enriched genes from the Human Protein Atlas^25^, we also found that 102 out of 134 orthogroups (which includes one-to-one, one-to-many, and many-to-many orthologs; see Methods) were conserved between humans and invertebrates. Strikingly, more than half of the 134 orthogroups (71) were conserved between humans and the sponge *Amphimedon queenslandica*–which lack obvious neurons and nervous tissue^38^–and 47 were conserved between humans and the closest non-animal outgroup, the choanoflagellate *Salpingoeca rosetta*. We also found that these genes are among the largest in each genome. This suggests that many of these large genes important for nervous systems have origins predating the diversification of animals and in many cases the emergence of dedicated neuronal cell types.

### Large ancient genes have gained the most isoforms

We observed that animals with expanded genomes have ancient, highly conserved genes that are acquiring new isoforms, mainly in larger genes (**Fig. 4**). Isoform numbers were obtained by quantifying annotated mRNAs for each gene and were compared for one-to-one orthologs. When we compared the set of large ancient genes among animals, we found that while orthologs of these genes are typically among the largest in each genome, they have become absolutely larger and more complex in vertebrates (**Fig. 1B**). This indicates that despite showing signs of strong purifying selection, surprisingly, these large ancient genes are acquiring many new sequences which may undergo positive selection and drive gene evolution.

**Figure 4.**
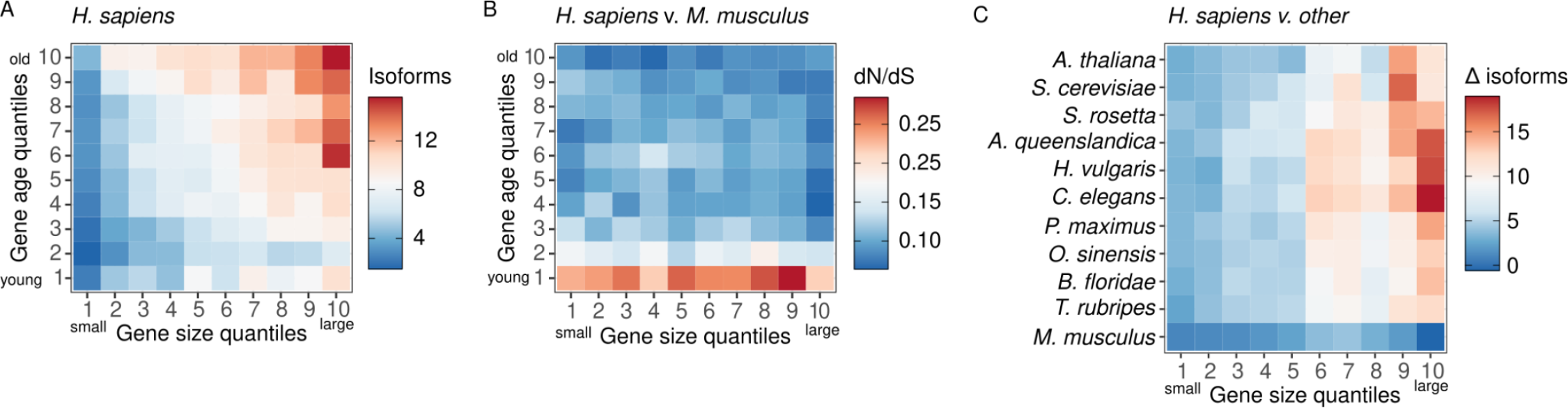
Large ancient genes have gained the most isoforms in humans. (**A**) Heatmap of human genes binned by gene size (10 bins) and gene age (10 bins). Heat colors show the average number of transcript isoforms per gene per bin. (**B**) Ensembl one-to-one orthologs between mouse and human showing average dN/dS scores as heat colors. (**C**) Heatmap of orthologs binned by human gene size (10 bins), with heat colors showing the average change in transcript isoforms between orthologs in human and each other species (e.g. *H. sapiens* - *A. thaliana, H. sapiens - S. cerevisiae*, etc.).

## Discussion

### Determinants of optimal gene size

By comparing the genomes and transcriptomes of diverse eukaryotes, we have outlined the contribution of gene size variation to the evolution of large neuronal genes. We propose the adaptive value in gene size expansion does not come from net gains directly, but rather in the addition of sites capable of sustaining beneficial mutations. Any change to the size of individual genes might disrupt coexpression dynamics. However, if these changes are balanced by the net changes in gene size of all coexpressed genes, coexpression might be maintained, while simultaneously generating the raw material for selection to act on. This could therefore effectively add new sites capable of sustaining beneficial mutations and potentiate gene architecture complexity in expanded genes. As the largest genes will have the largest absolute expansion of sequence space, these genes have the most potential to gain novel functions and expression patterns.

### Gene size and expression timing

Gene size directly affects expression timing and thus may contribute to the precise coordination of gene expression required by many biological processes. The effect of gene size on expression timing was first appreciated in lambda phage with the description of long, late operons^39^. When the size and abundance of introns in eukaryotic genes was discovered, these were likewise anticipated to have substantial effects on gene expression timing. This idea was articulated in the intron delay hypothesis, which postulates that intron size contributes to a time delay and aids the orchestration of gene expression patterns^40^. Several studies have since provided evidence that intron and gene size play a role in embryonic development by affecting transcriptional kinetics (see Swinburne and Silver 2008^41^ for a review). Additionally, highly expressed genes^29–31^ and genes involved in rapid stress response^32^ tend to have shorter introns, suggesting that selection for efficiency acts to reduce the time and energy costs of transcription. Together with our results, it appears that many biological processes involve genes with similar sizes, and that gene sizes may be evolving in part from selective pressure for expression timing.

### Gene size expansion and the addition of adaptive sites

The rate at which a gene under selection accrues beneficial substitutions is thought to be rapid at first, and eventually slows as the supply of sites capable of sustaining beneficial mutations (often referred to as “adaptive sites”) are depleted^42,43^. Under the “increasing constraint” model^44^, a newly born gene evolves under weak negative or positive selection, and later evolves primarily under strong negative selection. More recent evidence supports a variation of this idea, which is that young genes experience more variable dN/dS values than old genes^33^.

Our study provides evidence that gene size expansion in genes under high constraint (i.e. large and ancient genes under strong negative selection) can facilitate acquisition of sites capable of sustaining beneficial mutations in the form of new exons and regulatory regions. These new DNA sequences are likely under weaker constraint than the original sequences and can thus contribute to evolution. Many new exons arise from within introns and tend to be cassette exons that are rarely incorporated into final transcripts (i.e. they are spliced out)^45,46^. Similar to neo-functionalization of duplicated genes, because the original function is maintained by the major isoforms, the new isoforms are less constrained by negative selection^46^ and can thus contribute to adaptive evolution^45^. Thus, we speculate that gene size expansion may be one mechanism by which genes under high constraint can gain new raw material under weak constraint and contribute to the evolution of molecular diversity.

Previously, it has been argued that weaker constraint is unlikely to have contributed to the evolution of primate nervous systems because their complexity necessitates a greater precision in gene function^47^. Conversely, based on the results of this study, we speculate that this *weaker* constraint (through gene size expansion) may have set the conditions for the evolution of complex nervous systems by providing substrate for adaptive evolution.

### Gene size expansion and nervous system evolution

Gene size expansion has been hypothesized to facilitate the evolution of complex nervous systems^7,14,48^. This is in large part because most of the largest animal genes are multi-isoformic, enriched for expression in nervous tissue, and predominantly encode synaptic proteins underlying the precise wiring of the nervous system^7,12–15^. Additionally, large genes have been shown to contribute to the extensive molecular diversity and complexity of vertebrate brains^14^. However, because most studies of complex nervous systems have focused on vertebrates, it remains unclear if any such relationship arose from historical contingency. Did any invertebrate animals with complex nervous systems independently undergo gene size expansion?

While many vertebrates have large brains, as well as some of the largest genomes and gene sizes among animals^3,5,7^, there are outlier species among invertebrates, such as the cephalopods. Cephalopods have the largest invertebrate nervous systems and exhibit complex behaviors rivaling many vertebrates^21^. It has been more than 680 million years since cephalopods and vertebrates shared a last common ancestor^23^, which likely had a compact genome and gene sizes, as well as a simple nervous system^49^. Several chromosome-level genome assemblies for cephalopods have recently been published^50–52^, and in our analyses we found a striking expansion of gene sizes similar to that seen in the vertebrate lineage (**Fig. 1B**; **Fig. 5C**). The fact that many of these large, complex genes are enriched for neuronal expression and function across diverse animals is consistent with the hypothesis that gene size expansion contributed to the tremendous molecular diversity and complexity observed within nervous systems.

**Figure 5.**
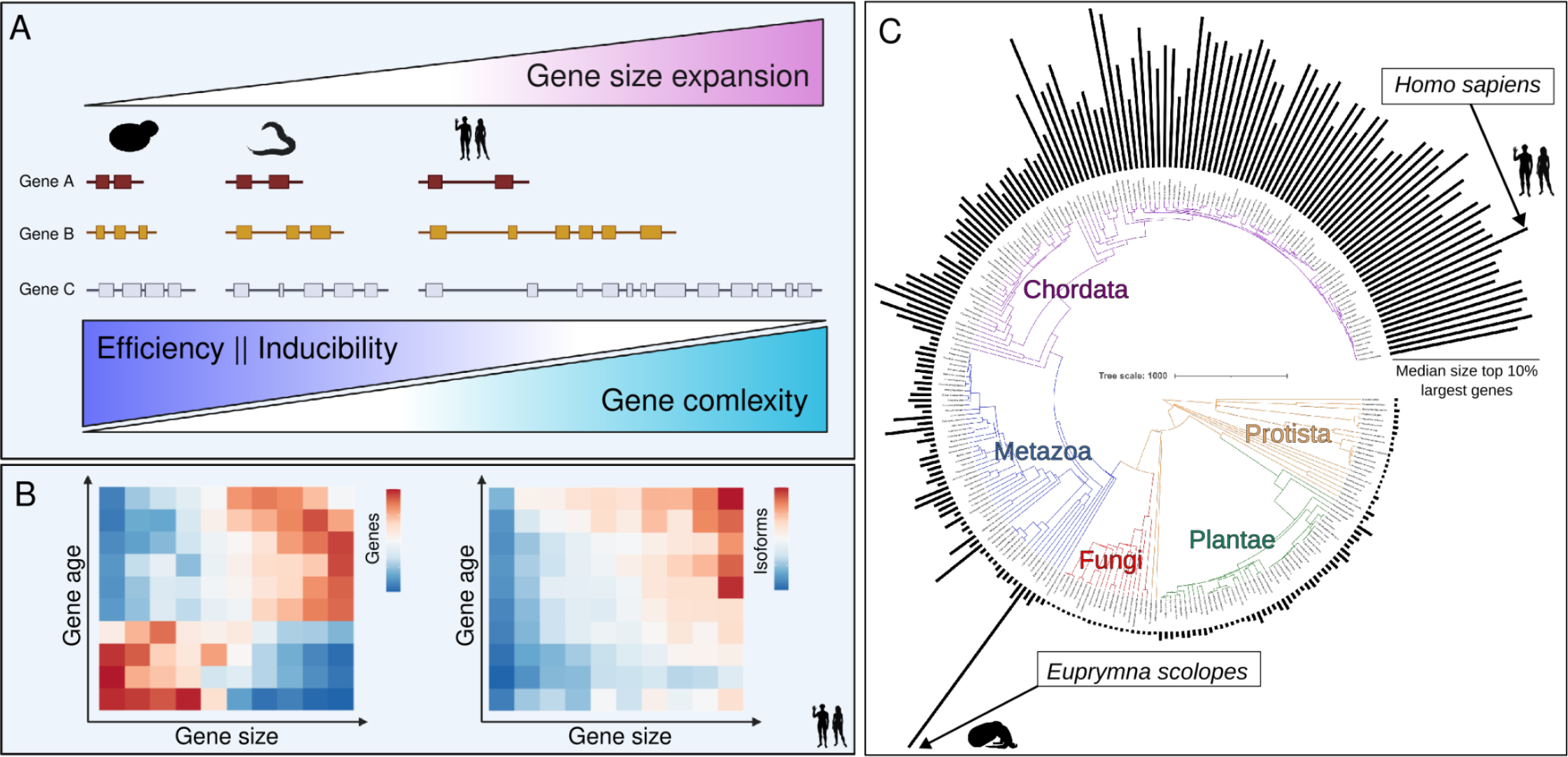
Model of gene size variation. (**A**) As genomes expand or contract, so does the intronic content of genes and hence gene sizes of eukaryotes. Larger genes are able to become more complex, but at the cost of inducibility and efficiency of expression. (**B**) Genes become large by being very old, and become complex by being large. (**C**) Species-specific differences in gene size variation may contribute to important differences in the potential for complex genes and phenotypes.

Of considerable interest in the context of models in which gene size expansion accompanies nervous system diversification are a number of counterexamples. For example, there are some animal genomes that underwent significant expansion (salamanders^53^, whale sharks^54^, lungfish^55^, grasshoppers^56^, etc.) without obvious increases in the complexity of their nervous systems relative to other animals. We speculate that gene size expansion is insufficient for gene architectural complexification, but may only set the conditions for further evolution by selection. It is also possible that the mechanisms by which genes and genomes expand impacts the mechanisms that generate novel regulatory elements and exons. For example, the diversity and composition of transposable element pools^57^ differs in a species-specific manner; more diverse transposable element pools may limit the acquisition of additional sequences by recombination, while some populations of transposable elements may be more or less likely to introduce regulatory modulation when inserted.

### Gene and genome size contraction

The focus of the current study was on gene and genome size expansion, but there are numerous examples of gene and genome size contraction as well. One example is the tomato russet mite, *Aculops lycopersici*, one of the smallest animals with the smallest known arthropod genome at 32.5 Mb^58^. There are few transposable elements (< 2% of the genome), small intergenic regions, and more than 80% of coding genes are intronless. Interestingly, 3’ introns were predominantly lost, which complements findings from other studies that 5’ introns are enriched for regulatory elements^59,60^. There are also cases of genome reduction among vertebrates, for example within the teleost fish, *Takifugu* (*T. rubripes*; 300 Mb)^61^.

If gene size expansion sets the conditions for added complexity, does that mean gene size contraction reduces the potential for complexity and adaptation? Future studies are needed to investigate these questions–in particular whether small genomes are evolutionary dead ends–which have implications for our understanding of how complex systems are generated or degenerated.

### Ancient events enabling recent adaptation

The genome design model^62^ posits that tissue-specific proteins have more complex architectures that explain the increase in their size. Extending this model, it has been argued that the complexity of large genes was already present at the base of the metazoan common ancestor^63^. Conversely, our results suggest that increases in the size of genes encoding tissue-specific proteins precede and potentiate the evolution of their more complex architecture. Rather than looking for the origins of gene-architectural and -regulatory complexity in the recent evolutionary history of any one species, our analysis suggests that ancient events established the necessary underlying conditions. The initial size of these genes may predispose them, over time, to becoming extremely large and accumulating sequences that selection can act on to generate complexity.

In conclusion, in this study we found that relative gene size is being maintained for most genes in each genome despite sometimes orders-of-magnitude changes in absolute gene sizes in orthologs among species. We found that most young genes are small, while virtually all larger genes are ancient. This includes the set of large neuronal genes, whose origins appear to predate the diversification of animals and in many cases the emergence of neurons and nervous systems. Maintaining relative gene size during evolution may facilitate the coordination of gene co-expression, while increases in absolute gene size may contribute to the evolution of novel gene structures and regulatory elements.

## Methods

### Gene and coding sequence sizes

Gene and coding sequence sizes in each species were obtained from Ensembl (ensembl.org)^64^. Gene start positions from the most 5’ exon were subtracted from gene end positions (+1) of the most 3’ exon to obtain a measure of gene size for each gene that excludes explicitly annotated 5′ and 3′ UTRs. Protein coding genes were selected using gene biotype information.

### Identification of orthologs

OrthoFinder^18^ was used to identify orthologs across several representative eukaryotes with chromosomal-level genome assemblies (excepting *S. rosetta* and *A. queenslandica*). OrthoFinder identifies groups of orthologous genes (orthogroups), which may include paralogs. The maximum size of all orthologs within each orthogroup was used. Ensembl was used for other “high-confidence”, one-to-one orthologs as indicated in the text.

### Gene ontology

*H. sapiens* gene ontology (GO terms) were obtained from Ensembl (ensembl.org)^64^, Ensembl genes 108, GRCh38.p13.

### Species phylogeny

Species phylogenies were obtained from TimeTree (timetree.org)^23^ and initially plotted using the Interactive Tree of Life^65^. Species outlines were obtained from phylopic.org.

### Gene ages

Gene ages were obtained from the GenOrigin database (genorigin.chenzxlab.cn)^37^. GenOrigin systematically infers gene age using a protein-family based pipeline (FBP) with Wagner parsimony algorithm, phylogeny derived from the TimeTree database (timetree.org)^23^, and orthology information from Ensembl Compara^22,66^.

### Species selection

The species analyzed in this study (**Table 1**) were chosen for the completeness of their genome assemblies, which has a significant impact on the quality and completeness of gene annotations. However, most complete genomes are biased for model organisms chosen for unique biological features with potential impacts on genome organization. As new genomes are sequenced to completion, the generality of these observations can be tested.

**Table 1.** Genome assemblies used.

| Species | RefSeq | Assembly | Level |
| --- | --- | --- | --- |
| *Homo sapiens* | GCF_009914755.1 |  | Chromosome |
| *Octopus sinensis* | GCF_006345805.1 |  | Chromosome |
| *Pecten maximus* | GCF_902652985.1 |  | Chromosome |
| *Hydra vulgaris* | GCF_022113875.1 |  | Chromosome |
| *Hormiphora californensis* | GCA_020137815.1 | Hcv1a1d20200309 | Chromosome |
| *Mus musculus* | GCF_000001635.27 |  | Chromosome |
| *Branchiostoma floridae* | GCF_000003815.2 |  | Chromosome |
| *Caenorhabditis elegans* | GCF_000002985.6 | WBcel235 | Chromosome |
| *Arabidopsis thaliana* | GCF_000001735.4 | TAIR10.1 | Chromosome |
| *Takifugu rubripes* | GCF_901000725.2 | fTakRub1.2 | Chromosome |
| *Saccharomyces cerevisiae* | GCF_000146045.2 | R64 | Chromosome |
| *Amphimedon queenslandica* | GCF_000090795.1 | v1.0 | Scaffold |
| *Salpingoeca rosetta* | GCF_000188695.1 | Proterospongia_sp_ATCC50818 | Scaffold |

### Statistical analyses

All statistical tests were performed in R version 3.5.0 (R Core Team 2018) and RStudio version 1.1.453 (RStudio Team 2015). All analyses will be made available as R scripts accompanied by data tables.

## Acknowledgements

Helpful input was provided by A. Herbert, D. Schultz, T. Rogers, O. Simakov, D. Lipman, K. Artiles, I. Zheludev, J. Chen, D. Galls, N. Hall, N. Jain, L. Wahba, M. Shoura, D. Jeong, U. Enam, E. Greenwald, and O. Ilbay. This work was funded by Stanford Wu Tsai Neurosciences Institute Interdisciplinary Scholar Fellowship (MJM),Stanford Genomics Training Program (5T32HG000044–22; MJM), Whitman Early Career Award (MJM), and NIH grant R35GM130366 (AZF).

## Extended Data

**Extended Data Figure 1.**
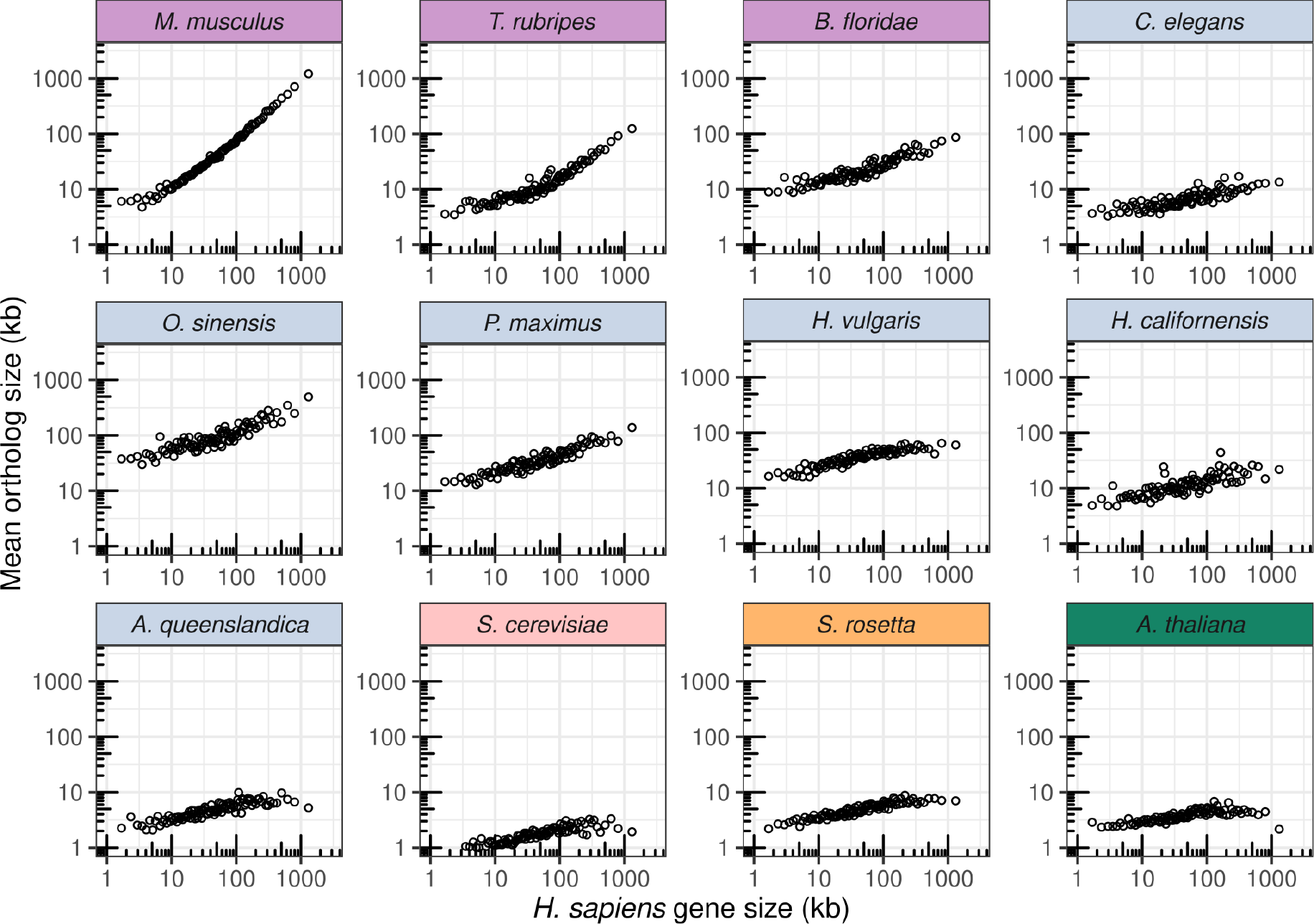
Scatter plots of ortholog gene size showing relative gene size preservation. For each group of orthologous genes between any two species, the max human gene size is shown versus the max ortholog size in other species. Each *H. sapiens* gene is binned into 50 quantiles, and the average gene size is shown for both *H. sapiens* genes and their orthologs for each bin. Box colors match clades: purple = vertebrates, blue = invertebrates, red = fungi, yellow = protists, green = plants.

